# OPTIMIZATION OF *RECOVERABLE CLEAR PLASTIC POT* (rCPP) METHOD AND VALIDATION OF MOLECULAR MARKS FOR THE SCREENING OF DEEP ROOTING OF RICE (*Oryza sativa* L.)

**DOI:** 10.1101/2024.07.14.603442

**Authors:** Mukhlis Syahril Pratama, Panjisakti Basunanda, Rani Agustina Wulandari, Muhammad Habib Widyawan

## Abstract

The generally of Rice plants have shallow roots and require large amounts of water for cultivation. However, water resources are becoming increasingly limited due to climate change. Therefore, efforts are needed to improve rice genetics in order to produce stable crop production in drought-prone areas. One strategy to avoid drought stress is to change the architecture of deeper rooting. The identification can be done by screening the root morphology. Most of the screening methods for deep rooting ability in rice are still *destructive*, so the proposed activity in this study is the optimization of the *Recoverable Clear Plastic Pot* (rCPP) method that can detect deep rooting ability in rice early and *non-destructively*. The method was further validated using InDel and SSR-based molecular markers designed based on the *DRO1* gene. The *DRO1* gene plays a role in increasing the root growth angle, so that the roots grow in a deeper vertical direction. The results showed that *Recoverable Clear Plastic Pot* (rCPP) optimization was able to show good screening ability for plant root characters *non-destructively*. Mayangan is the best drought-tolerant genotype based on deep rooting characters that have root distribution centered on the *inner circle*. The relationship between deep rooting screening using the DRO1 marker provides results that are in line with deep rooting screening through the *Recoverable Clear Plastic Pot* (rCPP) method.

**Highlight:**

1. This research is part of an effort to improve rice genetics in order to produce stable crop production in drought-prone areas.
2. The research activities used the optimization *Recoverable Clear Plastic Pot* (rCPP) method.
3. The rCPP method was validated using InDel and SSR-based molecular markers designed based on the *DRO1* gene.
4. Optimization of rCPP shows good screening ability for plant rooting characters in a *non-destructive manner*.

## Introduction

Conventional rice cultivation is done in paddy fields that require heavy water irrigation. Rice plants require 3000-5000 liters of water to produce one kilogram of rice (Pathak *et al*., 2011). Limited water availability for agricultural land can cause a decrease in rice production (Fuadi *et al*., 2016). According to BPS (2021), the productivity of paddy rice and field rice produced from rice cultivation with insufficient water availability is only about 43.45 ku/ha and 39.63 ku/ha, respectively. On the other hand, when water availability is sufficient, the average productivity of paddy rice can reach 52.07 ku/ha and the average productivity of field rice is 40.95 ku/ha. Rice plants generally have shallow roots so their cultivation requires large amounts of water. However, the current condition of water resources is increasingly limited due to climate change. Therefore, efforts are needed to improve rice genetics in order to produce stable crop production in drought-prone areas.

Drought is a limiting factor that can cause crop stress. Therefore, rice cultivars that can provide high yields under water shortage conditions need to be developed. Rice breeding programs for rainfed or upland rice conditions are directed towards the establishment of deep, dense and strong root systems (Mackill *et al*., 1996). Deep root systems are controlled by various genes that influence vertical and long root growth. Uga *et al*. (2011) reported that the *DRO1* gene in rice located on chromosome 9 is associated with deep rooting traits. *DRO1* is a gene that expresses deep rooting in rice plants. *DRO1* helps rice plants in utilizing soil water by increasing the angle of root growth so that it is tolerant to drought stress. In addition to *DRO1* which is related to the nature of deep rooting in rice, according to Yuka *et al*. (2011) found several genes that have the same properties include *DRO2* located on chromosome 4, *DRO3* on chromosome 7, and *DRO4* on chromosome 2.

In helping to phenotypically select rice plants that have deep rooting systems, plant breeders have developed various deep rooting screening methods; one of which was carried out by Ghozali and Supriyanta (2022) by developing the *Recovery Clear Pot Plastic Net* (rCPPN) method which is a combination of ideas from the *plastic net* and *clear pot methods* so that it has the ability to detect deep rooting quickly and easily in the seedling phase. However, the method in its implementation is still destructive so that the selected plants must be sacrificed. The development of this method can potentially be done *non-destructively so that the* selected plants can be allowed to grow back to maturity after observing deep rooting characters.

This research proposes an optimization of the rCPPN method by eliminating the use of *nets* and using one type of media and optimizing the capture of root images. This method is tentatively called *Recoverable Clear Plastic Pot* (rCPP). The main observation made in the rCPP method is to look at the angle of root growth based on the root photo image. This image is used to screen for deep rooting characteristics in a number of rice cultivars collected by the Department of Agricultural Cultivation UGM.

The results of the screening method were further validated using insertion and deletion (InDel) and *simple sequence repeats* (SSR) based molecular markers designed based on the *DRO1* gene. Several high-yielding rice cultivars with water stress resistance have been released, such as ‘Kasalath’ and ‘Bluebonnet’. Bluebonnet is an *indica* rice cultivar resulting from the crossing of Philippine drought-tolerant rice ‘Rexoro’ and ‘Fortuna’, which was released in Texas in 1944 (Tabien *et al*., 2008). Kasalath is a group of Aus rice, known to be tolerant to drought and high temperatures, and is often used as material in plant breeding research (Thomson et *al*., 2007; Fukuta *et al*., 2012). Gadjah Mada University has also produced strains of “amphibian rice” adaptive to dryland cultivation, referred to as the Gamagora series, totaling 10 strains. One of the Gamagora strains has been released under the name ‘Gamagora 7’ as a superior rice for paddy field cultivation. In this study, dryland-adapted rice was compared with paddy rice as a research material to validate the *Recoverable Clear Plastic Pot* (rCPP) method as a screening tool for deep rooting traits and at the same time link it to genetic markers derived from the *DRO1* gene.

## Materials and Methods Plant materials

The research was conducted for six months from July to December 2023 in Greenhouse and Laboratory of Plant Genetics and Breeding, Faculty of Agriculture, Universitas Gadjah Mada. The planting materials used in this study were 15 rice genotypes derived from Gamagora strains and the Faculty of Agriculture collection, namely Gamagora-2, Gamagora-4, Gamagora-5, Gamagora-7, Gamagora-8, Gamagora-10, Kasalath, Situ bagendit, Salumpikit, Bluebonnet, Inpago-12, Mayangan, Ciherang, M70D, and IR-64.

### Experimental design and agronomic practices

This research method used a randomized complete block design (RCBD) consisting of one factor with three blocks. Treatments consisted of 15 rice genotypes, with Ciherang as a negative control and Blubonnet as a positive control, so the number of experimental units was 45. Each experimental unit consisted of 3 plants. In this study, plants were placed under moderate stress conditions (60% field capacity) to trigger the formation of deep rooting ability. The 60% field capacity treatment was carried out by watering the plants with 30 ml of water every two days into the pots. The deep rooting screening method used refers to the method conducted by Ghozali and Supriyanta (2022) and Han *et al*. (2016) with several innovations, which include modifying the size of the pot, the position of planting rice in the pot, removing the *mesh net*, replacing the planting medium with sand, and observing the root architecture with panoramic photography at an angle of 120°. Rice was planted in the pot wall area after which the pot was placed in a plastic tub to be tilted at an angle of 30^0^ which aims to get a portrait of rice roots that are expected to stick to the pot wall as a whole. Roots that successfully penetrate or are in the *inner circle* are roots that are characterized by deep rooting (root growth angle of ≥ 60° measured from the soil surface to the bottom of the root). Making the *inner circle* area by looking at the results of root architecture photos, then drawing a straight line along 3.5 cm below the root growth point and pulling an angle of 30° to the right side and left side and pulling the line down (Figure 1).

**Figure 1.**
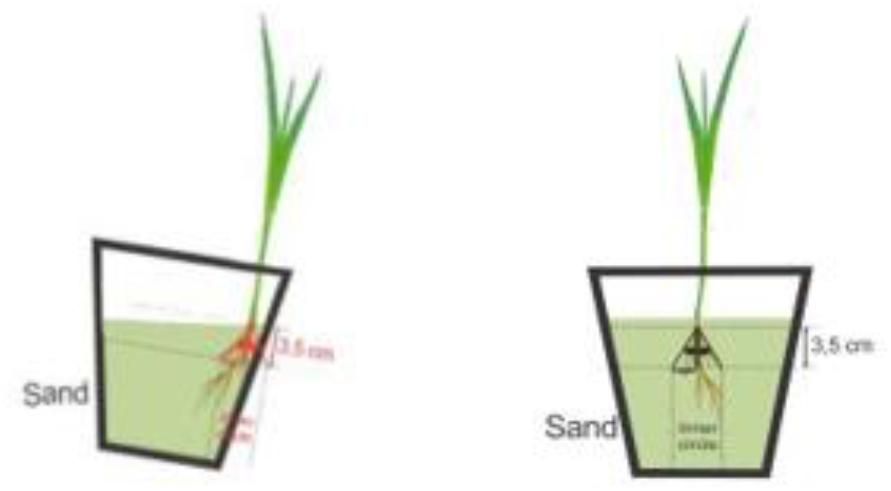
Pot position drawing of rCCP method

Fractal analysis was conducted using *box counting* method. With the help of *Corel Draw* and *Photoshop* applications, the results of panoramic photos of the roots were framed with rectilinear boxes each containing (r) 1; 2; 2.5; 4; 5; 10; 12.5; 20 cm (Tatsumi *et al*., 1995). The number of squares cut by the root as N(r) and recorded for each square containing r, then calculated the value of log N(r) and log r. The result of the equation will get log N (r) = -D log r + log K, where the value of D is the fractal dimension (FD) and log K is the fractal abundance (FA). In this study, fractal analysis was conducted in two ways, namely based on the total root architecture (FT) and the root architecture in the *inner cicle* (FI). FT is measured by looking at the entire root that cuts the box frame that has been determined, while FI is measured by looking at the root in the *inner circle* area that cuts the box frame. Furthermore, the ratio of the results of the FD and FA values in both analyses between FT and FI was made.

#### DNA isolation and amplification

DNA extraction was performed using the CTAB method (Doyle & Doyle, 1990). Leaf samples of two-week-old seedlings were ground using a mortar and pestle with liquid nitrogen. DNA quantification was performed using Nanodrop®. PCR amplification was carried out with two pairs of InDel primers namely *DRO-LIR* and *DRO-LKP* and two pairs of SSR primers namely RM 7424 and RM 24393 in Table 1. PCR amplification stages consist of *initial denaturation* at 95 ° C for 5 minutes, *denaturation* at 95 ° C for 1 minute, *annealing at* 55-56.9 ° C for 5 minutes, *extension at* 72 ° C for 1 minute 30 seconds, and *final extension at* 72 ° C for 7 minutes. The denaturation, *annealing*, and elongation stages were carried out for 36 cycles.

**Table 1.**
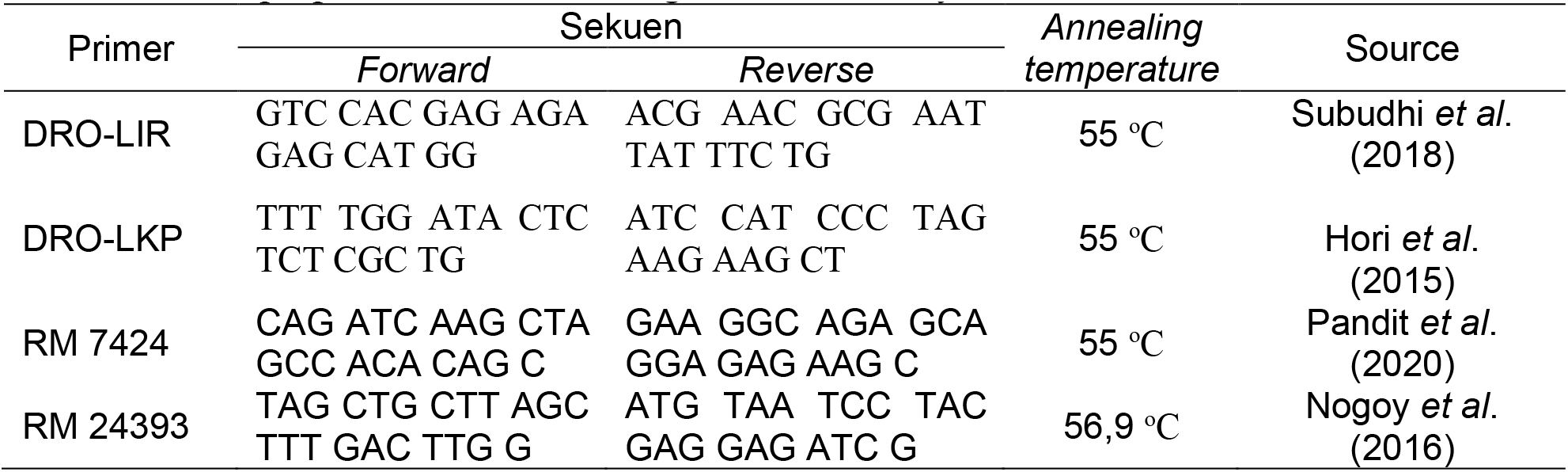
Primer properties of the inner ring used in the study.

### Statistical analysis

Data on morphological observations of 15 rice genotypes which included variables of plant height, root length, root diameter, and number of deep roots were analyzed using ANOVA at the α level of 5% while differences in treatment means were calculated by *Scoot-Knott* further test at the 5% level. Data from root panoramic photographs were analyzed using fractal analysis according to Tatsumi *et al*., 1989. using SAHN (*Sequential, Agglomerative, Hierarchical, and Nested*) analysis with UPGMA (*Unweight Pair-group Method Arithmatic Average)* method. The relationship between rice genotypes was analyzed using a *cluster heatmap* using a combination of morphological data and morphological data. The initial step of molecular data was first analyzed using Mahalanobis distance analysis. Mahalanobis distance (MD) is the distance between two points in multivariate space. The formula for calculating Mahalanobis distance is given as follows (Mahalanobis, 1936):

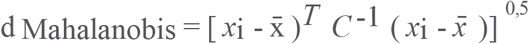

The mahalanobis distance results obtained single data in each geotype. Furthermore, the results of morphological and mahalanobis data are standardized to make the data uniform so that it can be visualized using a *cluster heatmap*.

## Results and discussion

### 1. Morphological Character Diversity

Most of the observation characters tested showed significant differences in the effect of the genotypes tested. Significant differences in all morphological characters tested indicate that the genetic diversity of the 15 genotypes is high, the results of the morphological characters of the 15 genotypes are presented in Table 2. The higher the genetic diversity, the higher the opportunity to obtain gene sources for characters to be improved (Boer, 2011).

**Table 2:** Morphological characters of 15 rice genotypes.

| Genotype | TT (cm) | PA (cm) | DA (cm) | JPD |
| --- | --- | --- | --- | --- |
| Gamagora-2 | 28.222 c | 8.744 b | 0.153 b | 1.926 b |
| Gamagora-4 | 27.000 d | 9.889 b | 0.16 b | 1.893 b |
| Gamagora-5 | 35.000 b | 8.733 b | 0.16 b | 2.09 b |
| Gamagora-7 | 36.777 a | 10.289 a | 0.173 b | 2.241 a |
| Gamagora-8 | 34,667 b | 8.578 b | 0.18 b | 1.912 b |
| Gamagora-10 | 34,111 b | 8.022 b | 0.167 b | 2.138 b |
| Kasalath | 38.667 a | 8.700 b | 0.173 b | 1.901 b |
| Situ Bagendit | 33.222 b | 9.367 c | 0.153 b | 1.936 b |
| Salumpikit | 37.112 a | 11.400 a | 0.22 a | 2.295 a |
| Bluebonnet (positive control) | 36.000 a | 11.333 a | 0.233 a | 2.263 a |
| Inpago-12 | 33.889 b | 10.289 a | 0.16 b | 1.97 b |
| Mayangan | 38.778 a | 11.833 a | 0.24 a | 2.295 a |
| Ciherang | 28.889 c | 6.500 c | 0.133 b | 1.825 c |
| M-70D | 33.000 b | 9.777 b | 0.147 b | 1.913 b |
| IR-64 (negative control) | 24.889 d | 4.545 d | 0.14 b | 1.803 b |

Notes: Means followed by the same letter in each column show no significant difference in the Scoot-Knott test at the 5% level, TT = plant height, PA = root length, DA = root diameter, and JPD = amount deep roots.

The results of plant height characters showed that the Mayangan genotype had a greater mean plant height than the other genotypes at 38.788 cm, but was as good as the positive control genotype, Bluebonnet, and significantly different from the negative control genotype, IR-64. The genotype with the lowest average plant height is IR-64 which is significantly different from other genotypes at 24.889 cm. These results are in accordance with Singh *et al*. (2018) that plants that are increasingly stressed by drought will interfere with plant height growth because cell turgor pressure decreases which causes cell metabolism to decrease so that the process of cell division and cell elongation will be inhibited. The results of the average root diameter showed that the Mayangan genotype had a thicker average root diameter of 0.24 cm, which was significantly different from the negative control genotype, IR-64, but as good as the positive control genotype, Blubonnet. The genotype with the lowest average root diameter and not significantly different from the negative control genotype was the Ciherang genotype with an average root diameter of 0.133 cm. This is in accordance with Yin *et al* (2017) that drought stress can increase the number of transport vessels and the diameter of transport vessels in the roots so that it helps in maximizing the absorption of nutrients and water in the soil.

Root diameter has an important role in the development and function of roots in finding water and nutrients. The results of the average root diameter in Table 2 can be seen that the Mayangan genotype has a thicker average root diameter of 0.24 cm, which is significantly different from the negative control genotype, IR-64, but as good as the positive control genotype, Bluebonnet. The genotype with the lowest average root diameter and not significantly different from the negative control genotype was the Ciherang genotype with an average root diameter of 0.133 cm. This is in accordance with Yin *et al* (2017) that drought stress can increase the number of transport vessels and the diameter of transport vessels in the roots so that it helps in maximizing the absorption of nutrients and water in the soil. The number of deep roots is the number of roots that are able to penetrate the *inner circle* that has been determined based on the method used. The results of the number of deep roots in Table 2 can be seen that the Mayangan and Salumpikit genotypes have a significantly different average number of deep roots and the largest from the negative control genotype of 2.295. Plant avoidance in dealing with drought stress is one of them by modifying deep rooting. The genotype with the lowest number of deep roots and significantly different from other genotypes is the Ciherang genotype at 1.825. The amount of deep rooting reflects roots that have a growth angle of 60° measured from the soil surface until the root grows (Han *et al*., 2016). According to Uga *et al*. (2013), deep rooting is a plant strategy to avoid drought stress so that it can increase the plant’s ability to extract water from deeper soil layers. The root growth angle expressed by the *DRO-1* gene is negatively regulated by auxin and is involved in cell elongation at the root tip which causes asymmetric root growth and vertical downward direction.

Based on the fractal values in Table 3, it is found that the genotypes Salumpikit, Bluebonnet, and Mayangan have fewer FD values in the FT/FI ratio than the other genotypes. This indicates that the branching of lateral roots in these genotypes has a narrower branching angle so that the intersection of the box that fills the fractal space does not differ much between the total fractal and the *inner* fractal. According to Tatsumi 1995, the fractal value is not determined by the length or area of the root system but is closely related to the branching sequence or density of the lateral roots. Therefore FD values may indicate changes in root branching patterns. In addition, lateral roots were found to be responsible for the major root part of the rice root system especially during drought conditions based on the results of Wang *et al* (2009).

**Table 3.**
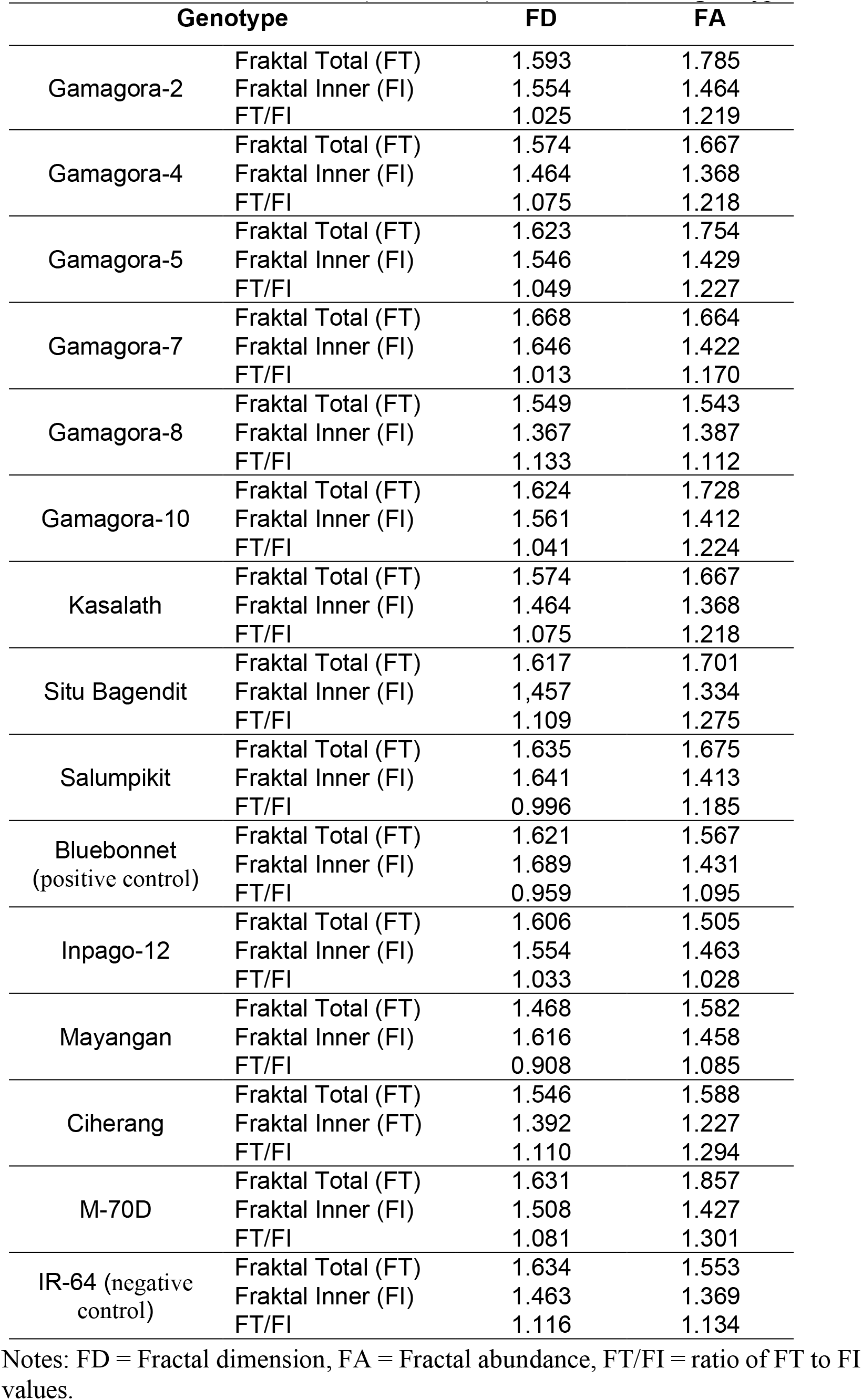
Fractal dimensions (FD and FA) of roots of 15 rice genotypes.

| Genotype |  | FD | FA |
| --- | --- | --- | --- |
| Gamagora-2 | Fraktal Total (FT) | 1.593 | 1.785 |
|  | Fraktal Inner (FI) | 1.554 | 1.464 |
|  | FT/FI | 1.025 | 1.219 |
| Gamagora-4 | Fraktal Total (FT) | 1.574 | 1.667 |
|  | Fraktal Inner (FI) | 1.464 | 1.368 |
|  | FT/FI | 1.075 | 1.218 |
| Gamagora-5 | Fraktal Total (FT) | 1.623 | 1.754 |
|  | Fraktal Inner (FI) | 1.546 | 1.429 |
|  | FT/FI | 1.049 | 1.227 |
| Gamagora-7 | Fraktal Total (FT) | 1.668 | 1.664 |
|  | Fraktal Inner (FI) | 1.646 | 1.422 |
|  | FT/FI | 1.013 | 1.170 |
| Gamagora-8 | Fraktal Total (FT) | 1.549 | 1.543 |
|  | Fraktal Inner (FI) | 1.367 | 1.387 |
|  | FT/FI | 1.133 | 1.112 |
| Gamagora-10 | Fraktal Total (FT) | 1.624 | 1.728 |
|  | Fraktal Inner (FI) | 1.561 | 1.412 |
|  | FT/FI | 1.041 | 1.224 |
| Kasalath | Fraktal Total (FT) | 1.574 | 1.667 |
|  | Fraktal Inner (FI) | 1.464 | 1.368 |
|  | FT/FI | 1.075 | 1.218 |
| Situ Bagendit | Fraktal Total (FT) | 1.617 | 1.701 |
|  | Fraktal Inner (FI) | 1.457 | 1.334 |
|  | FT/FI | 1.109 | 1.275 |
| Salumpikit | Fraktal Total (FT) | 1.635 | 1.675 |
|  | Fraktal Inner (FI) | 1.641 | 1.413 |
|  | FT/FI | 0.996 | 1.185 |
| Bluebonnet<br>(positive control) | Fraktal Total (FT) | 1.621 | 1.567 |
|  | Fraktal Inner (FI) | 1.689 | 1.431 |
|  | FT/FI | 0.959 | 1.095 |
| Inpago-12 | Fraktal Total (FT) | 1.606 | 1.505 |
|  | Fraktal Inner (FI) | 1.554 | 1.463 |
|  | FT/FI | 1.033 | 1.028 |
| Mayangan | Fraktal Total (FT) | 1.468 | 1.582 |
|  | Fraktal Inner (FI) | 1.616 | 1.458 |
|  | FT/FI | 0.908 | 1.085 |
| Ciherang | Fraktal Total (FT) | 1.546 | 1.588 |
|  | Fraktal Inner (FI) | 1.392 | 1.227 |
|  | FT/FI | 1.110 | 1.294 |
| M-70D | Fraktal Total (FT) | 1.631 | 1.857 |
|  | Fraktal Inner (FI) | 1.508 | 1.427 |
|  | FT/FI | 1.081 | 1.301 |
| IR-64 (negative<br>control) | Fraktal Total (FT) | 1.634 | 1.553 |
|  | Fraktal Inner (FI) | 1.463 | 1.369 |
|  | FT/FI | 1.116 | 1.134 |

Notes: FD = Fractal dimension, FA = Fractal abundance, FT/FI = ratio of FT to FI values.

Fractal analysis based on *inner circle* (FI) was conducted because research in general about fractal analysis on plant roots, only seen from how much root branching is produced and how many roots fill the volume of soil explored. This is less effective in distinguishing drought-resistant genotypes from those that are not resistant, because not necessarily the roots with many branches grow in the vertical direction so that they are unable to penetrate the deep soil to absorb water in a drought-stressed situation. Genotypes M-70D and Ciherang have FA values in the FT/FI ratio greater than the other genotypes, this is because the branching of the roots is very spread out and when viewed through the fractal *inner* genotypes only a few that cut the *inner* area. According to Wang *et al* (2009), FA is closely related to the volume of soil explored by the root system, which has been proposed by other studies (Tatsumi *et al*., 1995; Tatsumi, 2001).

### 2. Molecular Character Diversity

In the amplification of genomic DNA from 15 rice genotypes, using two insertion and deletion (InDel) primers and two SSR primers, all polymorphic primers were found. Of the total primers amplified, the average polymorphic fragment per primer was 5.438. Dendrograms based on UPGMA analysis grouped the 15 genotypes into distinct groups. Jaccard coefficient similarity ranged from 0.13 to 1.00 (Figure 2).

**Figure 2.**
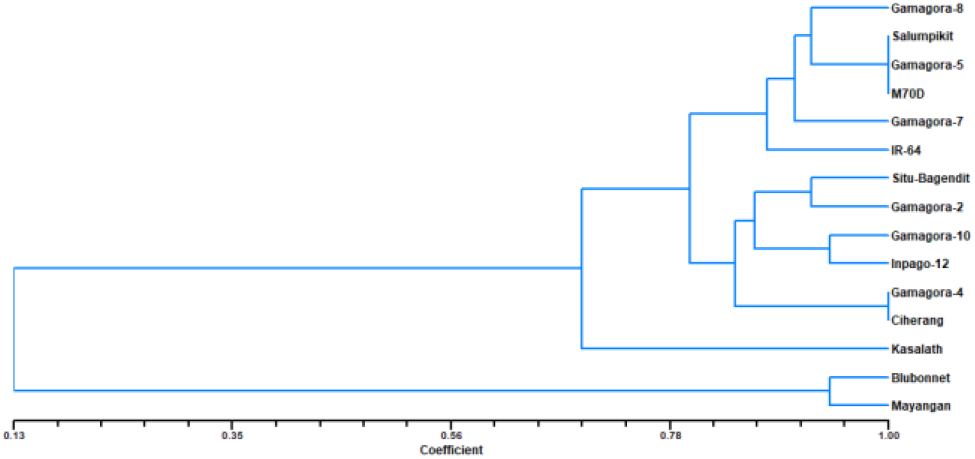
*Clustering* dendrogram of molecular analysis of rooting characters in 15 rice genotypes

At 13% genetic similarity the genotypes were grouped into 2 clusters. Cluster I consists of Blubonnet and Mayangan genotypes, Blubbonet is a positive control genotype that has the *DRO-1* gene that can express deep-rooting characters, so the genotypes in cluster I are genotypes that have deep-rooting characters. Cluster II is divided into three subclusters at a genetic similarity of 80.2%, namely subcluster I consisting of the genotypes Gamagora-8, Salumpikit, Gamagora-5, M70D, Gamagora-7, and IR 64. Subcluster II consists of the genotypes Situ Bagendit, Gamagora-2, Gamagora-10, Inpago-12, Gamagora-4, and Ciherang. Subcluster III consists of the Kasalath genotype. Subcluster I contained the IR-64 genotype which is a negative control genotype so that the genotypes in subcluster I may not have the *DRO-1* gene. Subcluster II has a similarity with subcluster I of 0.055 which means that most of the genotypes in subcluster II do not have the *DRO-1* gene. Subcluster III has a similarity with subcluster I of 0.275 which means that the genotypes in subcluster III mostly do not have the *DRO-1 gene*.

### 3. Correlation of Morphological and Molecular Results

Based on the data from the morphological analysis, it was found that the Mayangan and Salumpikit genotypes were not significantly different from the positive control genotype Bluebonnet in the characters of plant height, root length, root diameter, and number of *inner* roots, which means that these genotypes have deep rooting characters that can help plants maintain relatively high tissue water potential despite water shortages. Molecular results data support that the Mayangan genotype is similar to the Bluebonnet positive control genotype which has the *DRO-1* gene that expresses deep rooting traits. Whereas the Salumpikit genotype has a very large coefficient of dissimilarity with the positive control genotype based on the molecular data generated, although the morphological results are not significantly different from the positive control genotype. This could be due to the fact that the morphological expression produced is more influenced by the environment or other resistance genes that can express deep rooting traits.

Based on mahalanobis analysis and standardization test of morphological and molecular data of 15 rice genotypes can be visualized using *Heatmap clustrer* (Figure 3). There are 2 clusters that are divided, in *cluster* I there is a positive control genotype, namely Blubonnet and in cluster II there is a positive control genotype, namely IR-62. So it can be concluded that *cluster* I consisting of genotypes Blubonnet, Salumpikit, Gamagora-7, Kasalath, Gamagora-10, Gamagora-5, Inpago-12, and Mayangan are drought-resistant genotypes based on deep rooting and leaf rolling characters, while *cluster* II containing genotypes IR-64, Ciherang, M70D, Gamagora-4, Gamagora-2, Situ Bagendit, and Gamagora-8 are genotypes that are not drought-resistant based on deep rooting and leaf rolling characters. Cluster I can be divided into three *subclusters. Subcluster* I consists of genotypes Blubonnet, Salumpikit, Gamagora-7 which have similarities in leaf drying characters, plant *recovery*, and *inner* fractal abundance. Subcluster II consists of genotypes Kasalath, Gamagora-10, Gamagora-5, Inpago-12 which have similarities in leaf drying characters. Subcluster III consists of the Mayangan genotype which has the best drought resistance results among other genotypes based on deep rooting and leaf rolling characters.

**Figure 3.**
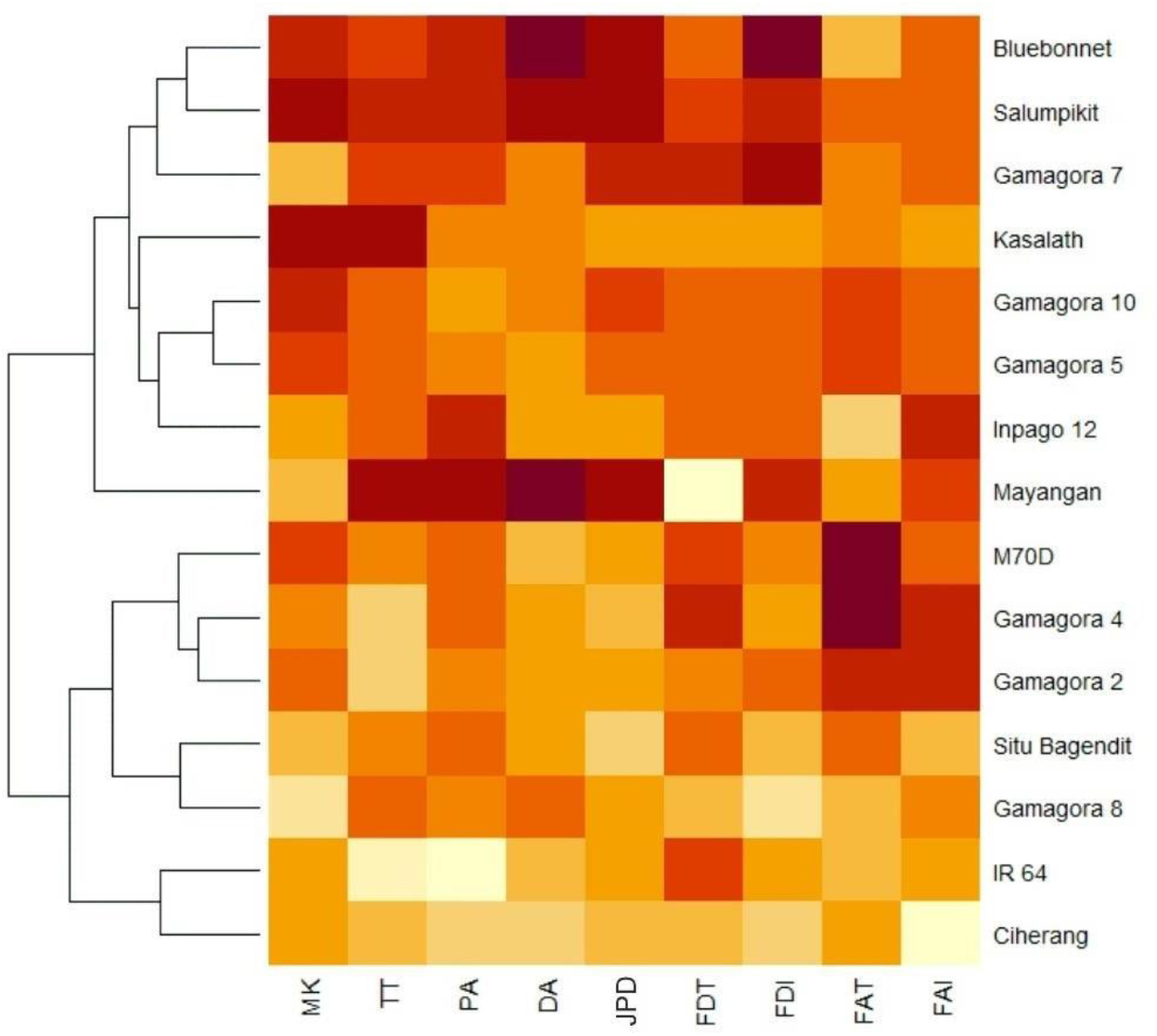
*Heatmap of clusters of* 15 rice genotypes

Optimization of the *Recoverable Clear Plastic Pot* (rCPP) method was able to show good screening ability for plant root characters *non-destructively*. The distribution of tolerant rice roots centered on the *inner circle* and the distribution of sensitive rice roots spread outside the *inner circle* (Figure 4) Mayangan is the best drought-tolerant genotype based on deep rooting characters that can help increase the angle of root growth so that plant roots grow vertically and can penetrate the subsoil to obtain water. The relationship between deep rooting screening using the *DRO1* marker gives results that are in line with deep rooting screening through the *Recoverable Clear Plastic Pot* (rCPP) method.

**Figure 4.**
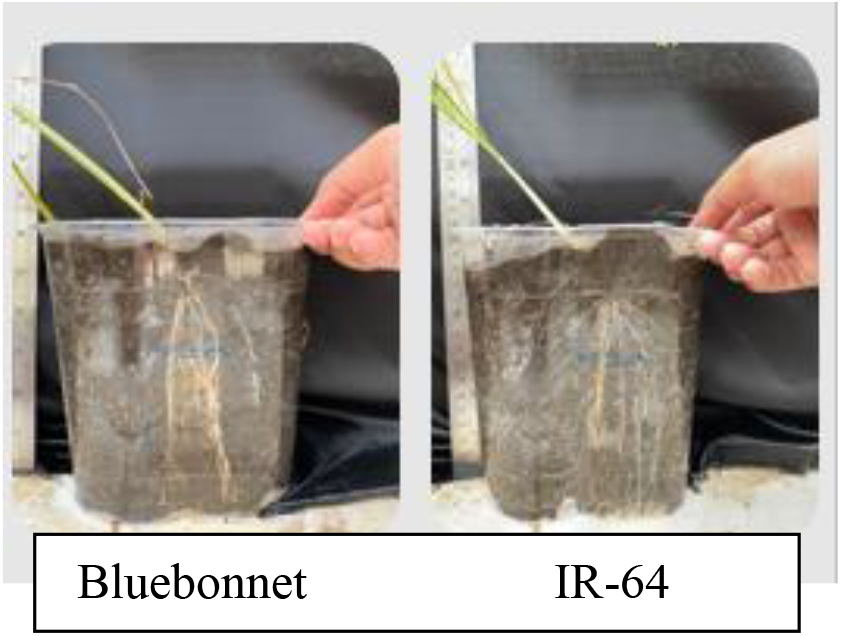
*Recoverable Clear Plastic Pot* (rCPP)

## Acknowledgements

Praise and gratitude to Allah SWT who has bestowed all His grace and guidance, so that the author can complete the writing of this scientific work. Thank you to my parents, Mr. Dr. Panjisakti Basunanda, S.P., M.P., 2. Mrs. Rani Agustina Wulandari, S.P., M.P., Ph.D, who have provided guidance, direction, criticism, advice and support.

## Author contributions

The author contributed to the preparation of the research proposal, the search and collection of data in the field, testing in the laboratory, and writing the research results.

---

**Gambar 1.**
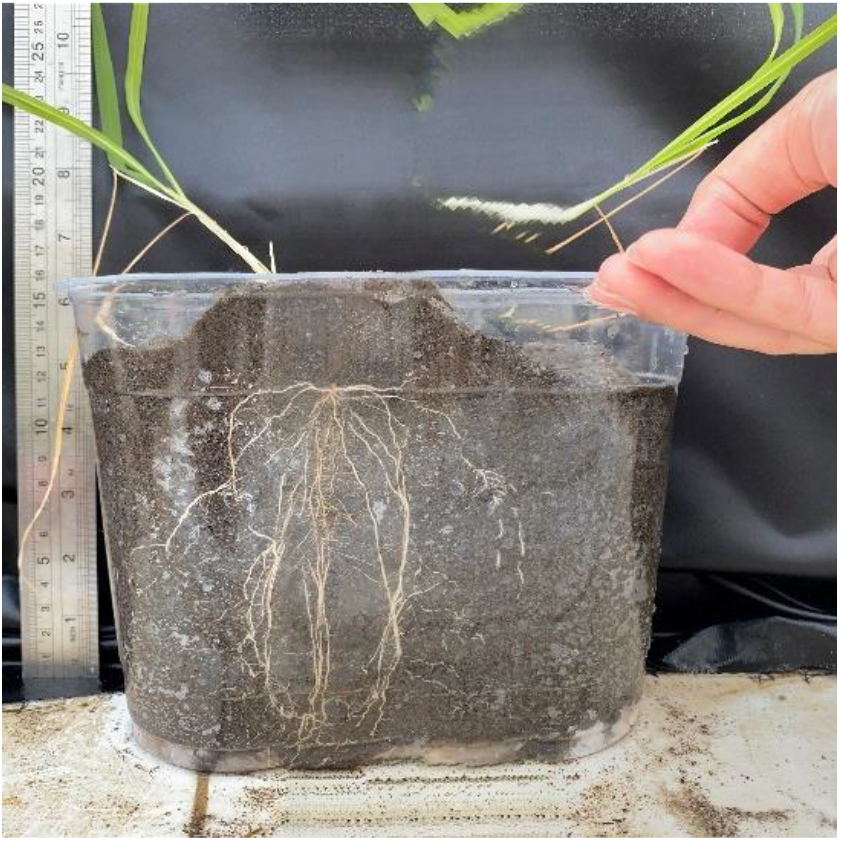
Hasil foto panorama genotipe Gamagora-2

**Gambar 2.**
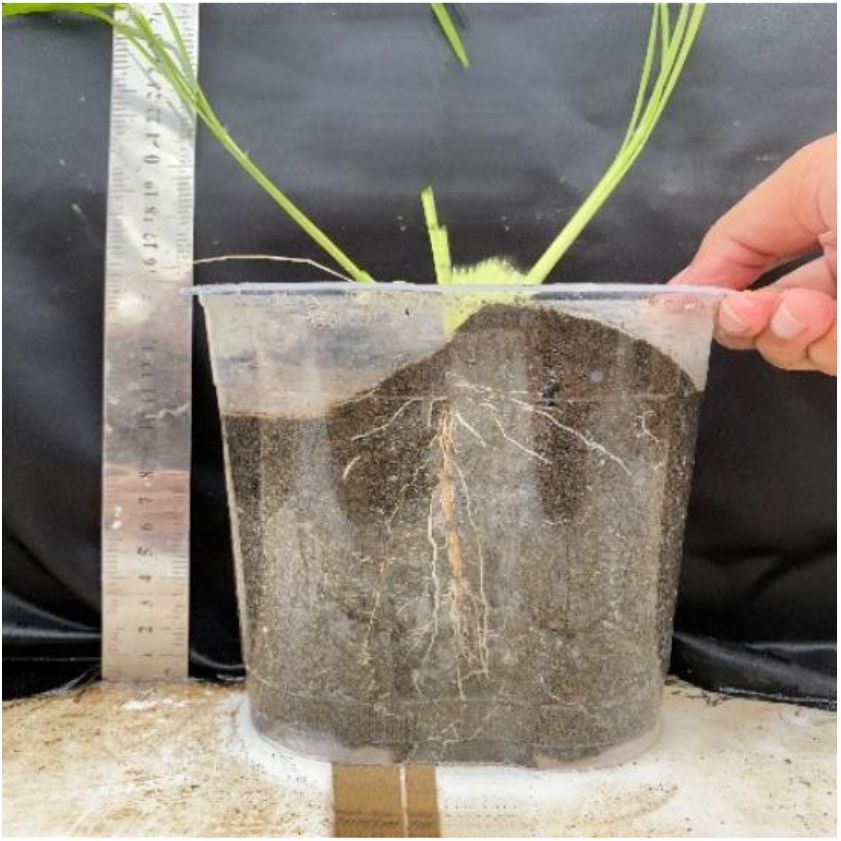
Hasil foto panorama genotipe Gamagora-4

**Gambar 3.**
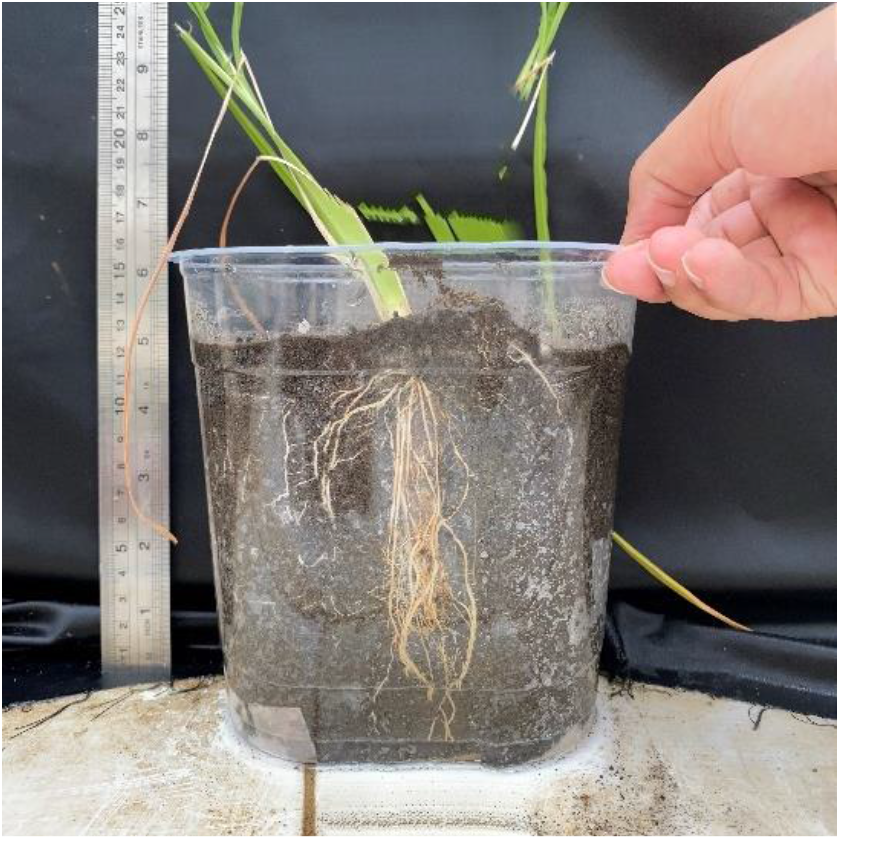
Hasil foto panorama genotipe Gamagora-5

**Gambar 4.**
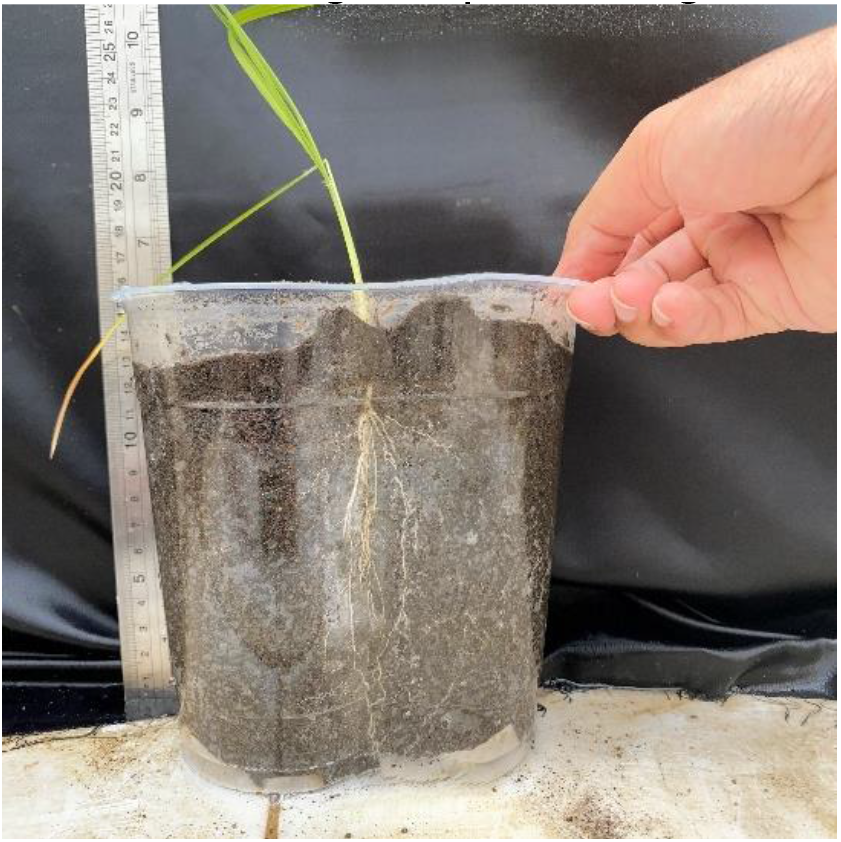
Hasil foto panorama genotipe Gamagora-7

**Gambar 5.**
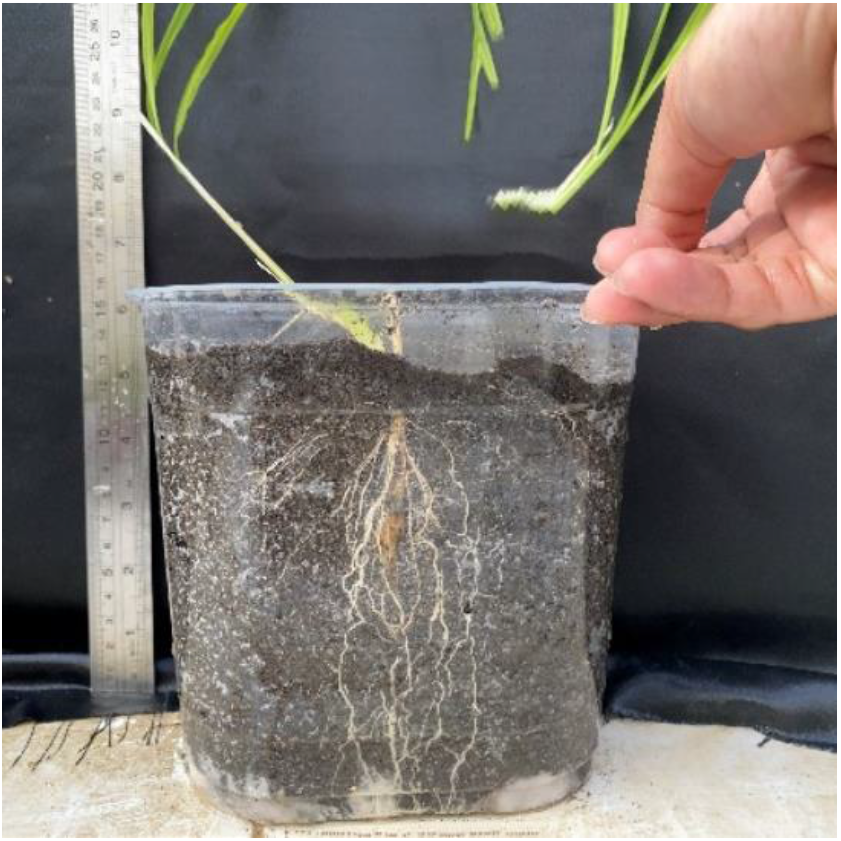
Hasil foto panorama genotipe Gamagora-8

**Gambar 6.**
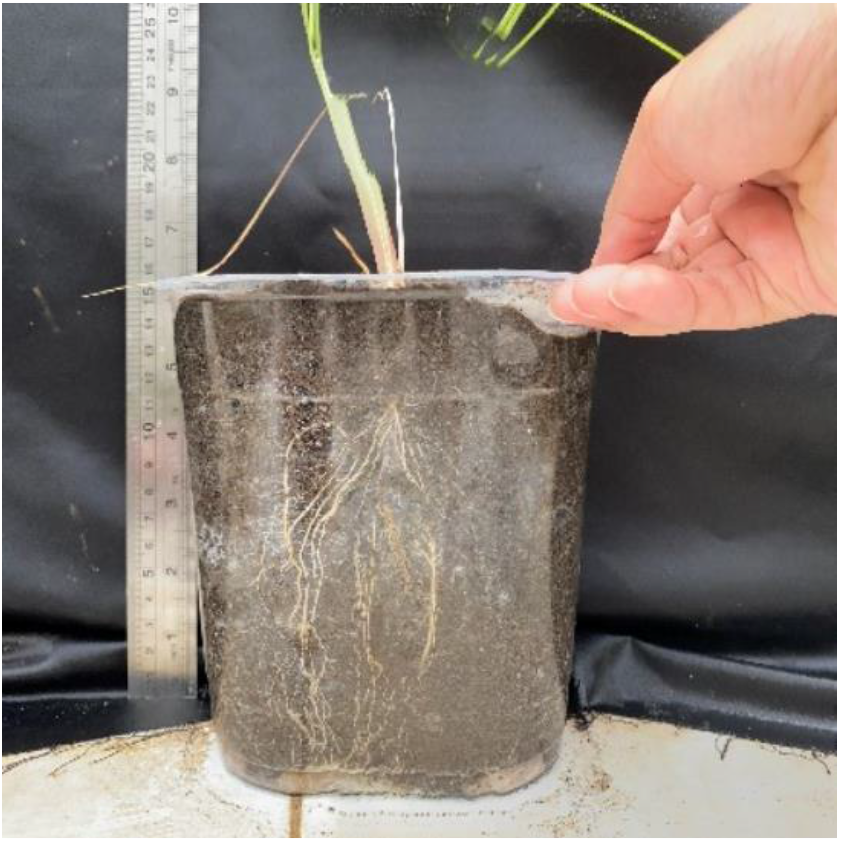
Hasil foto panorama genotipe Gamagora-10

**Gambar 7.**
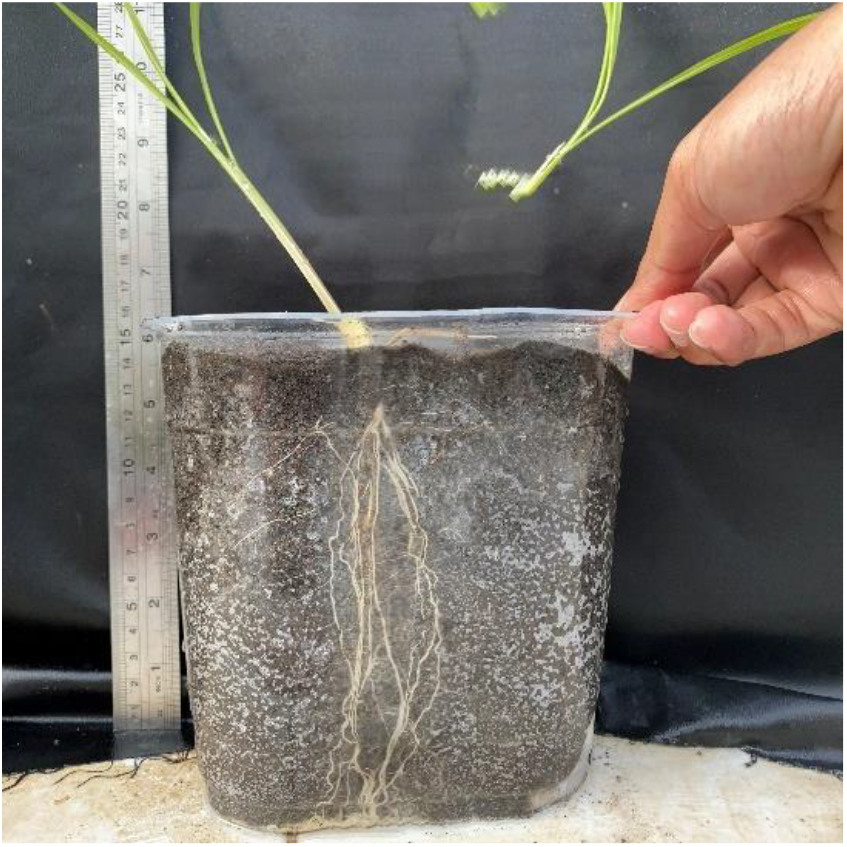
Hasil foto panorama genotipe Kasalath

**Gambar 8.**
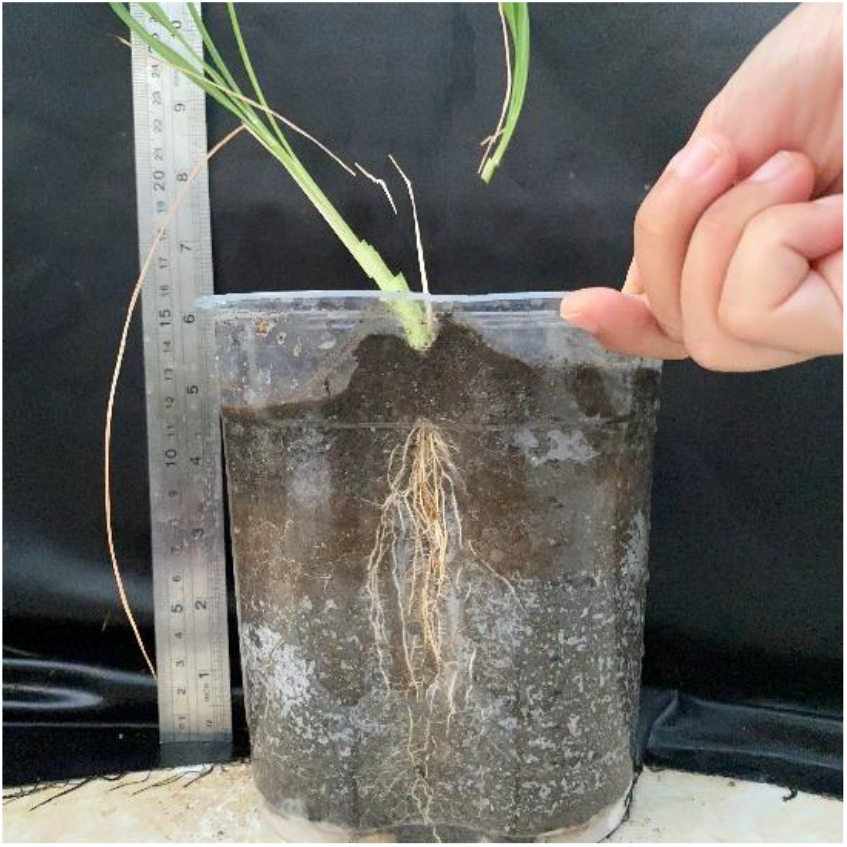
Hasil foto panorama genotipe Situ Bagendit

**Gambar 9.**
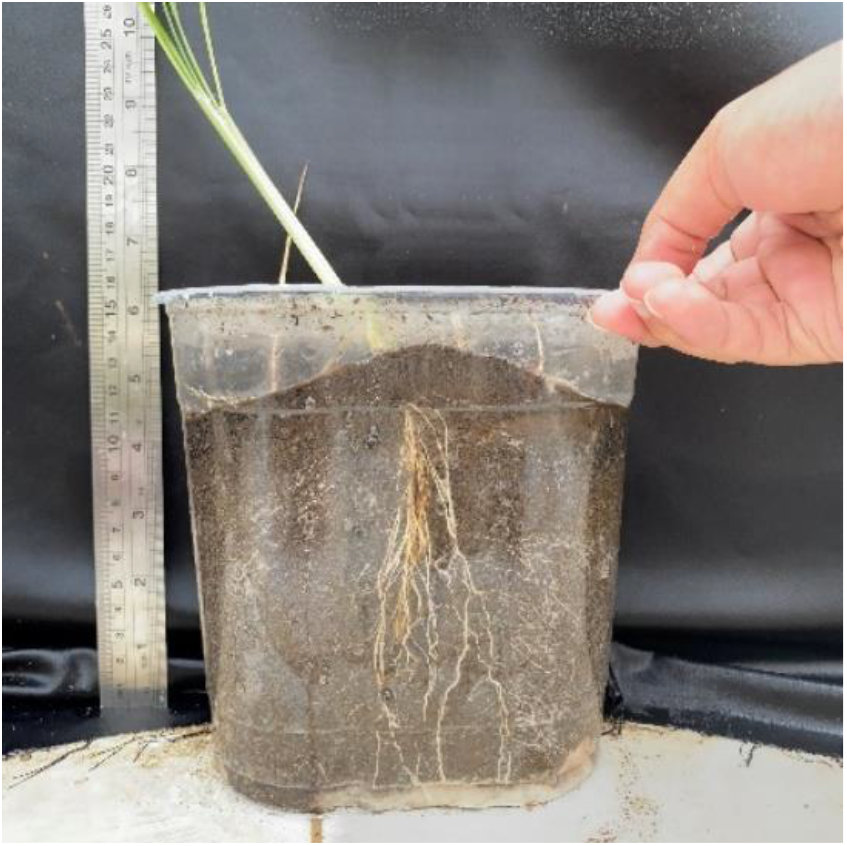
Hasil foto panorama genotipe Salumpikit

**Gambar 10.**
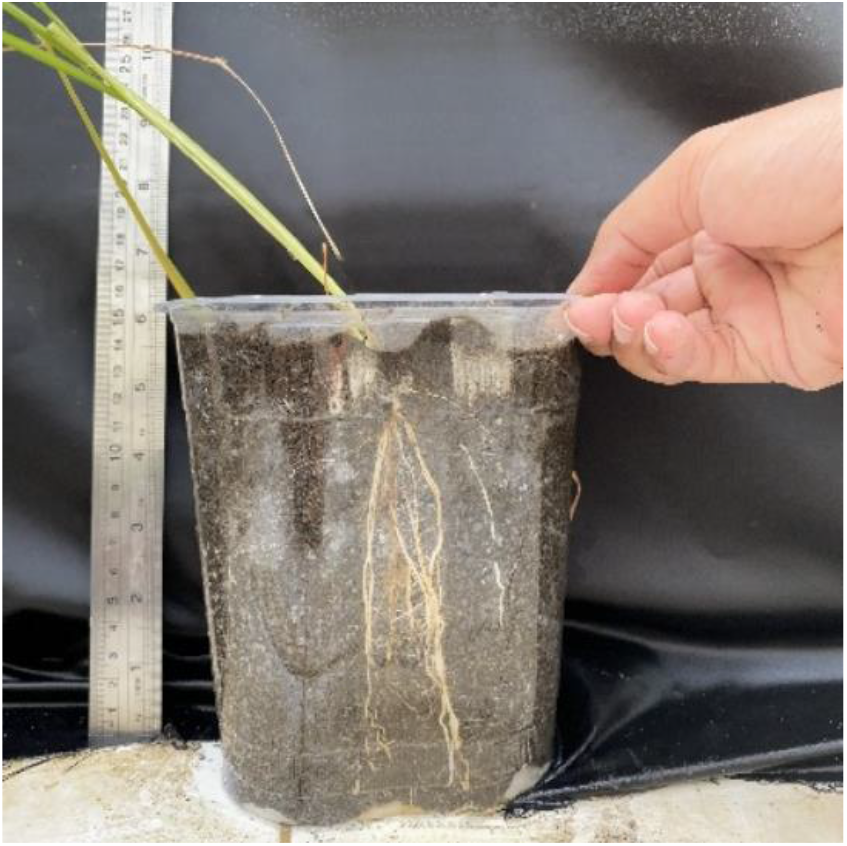
Hasil foto panorama genotipe Bluebonnet

**Gambar 11.**
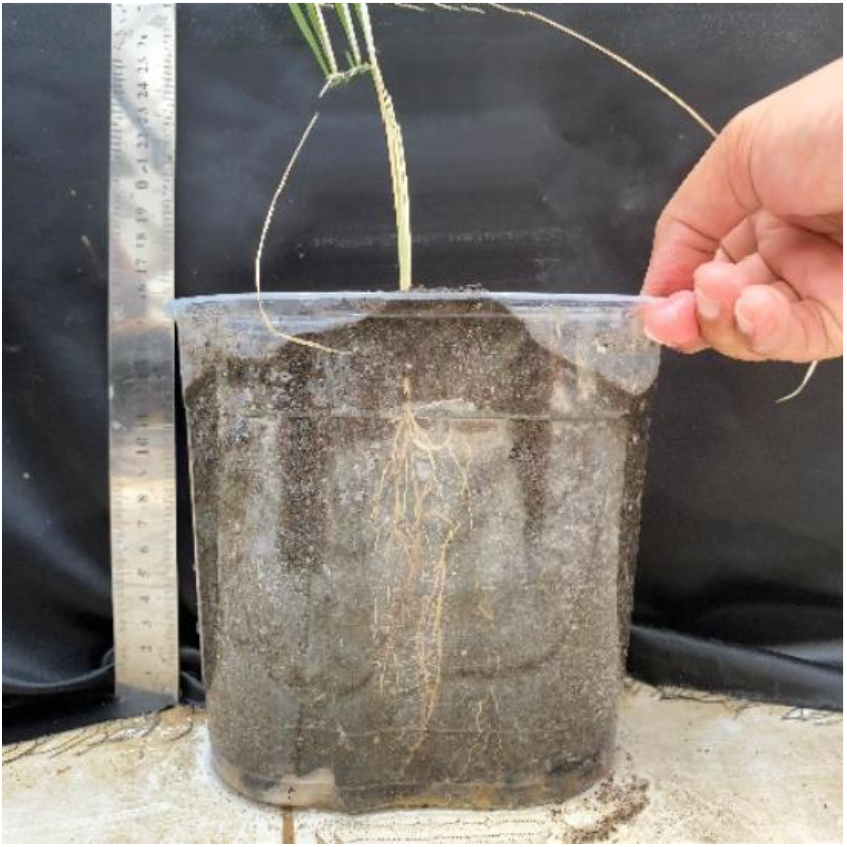
Hasil foto panorama genotipe Inpago-12

**Gambar 12.**
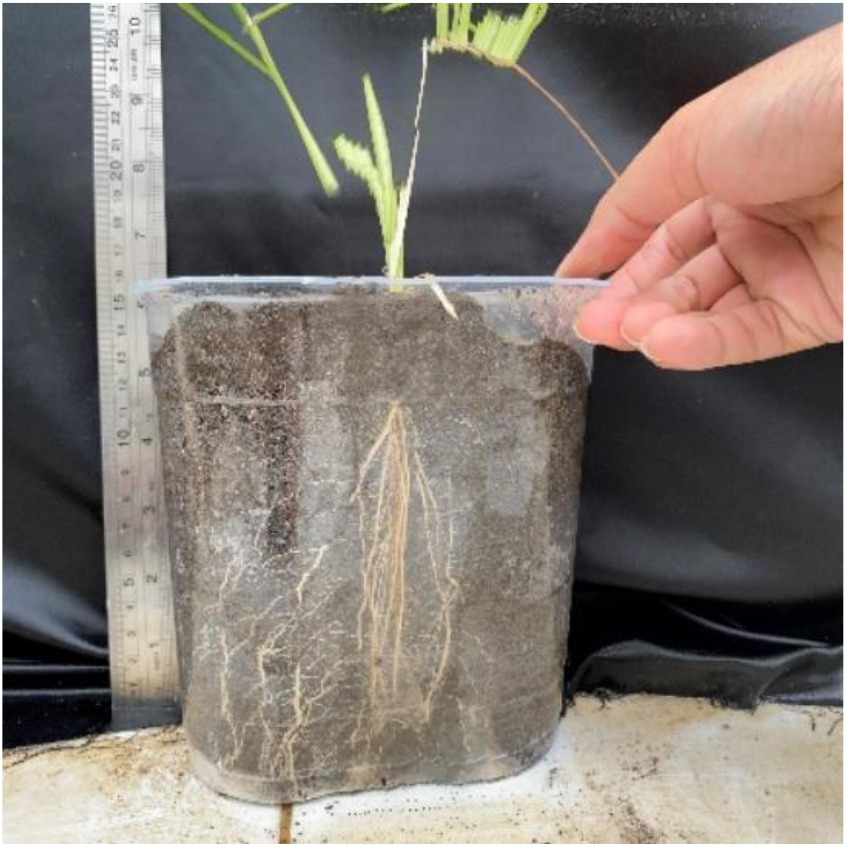
Hasil foto panorama genotipe Ciherang

**Gambar 13.**
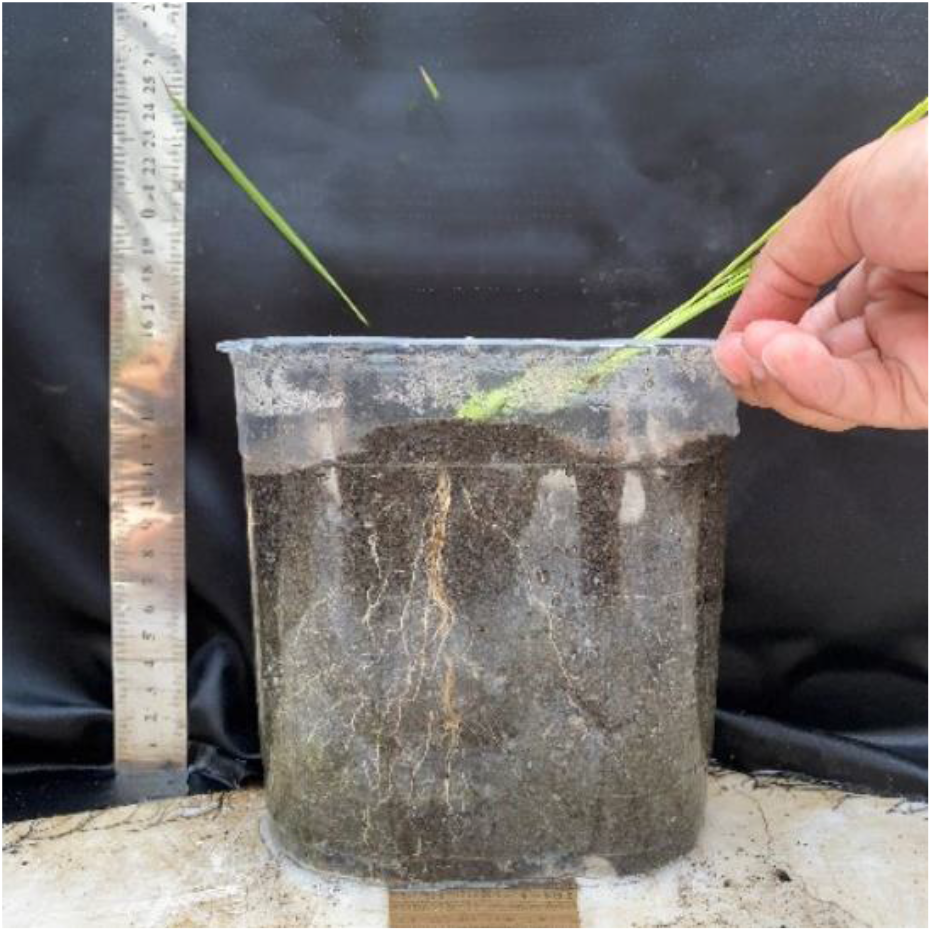
Hasil foto panorama genotipe Ciherang

**Gambar 14.**
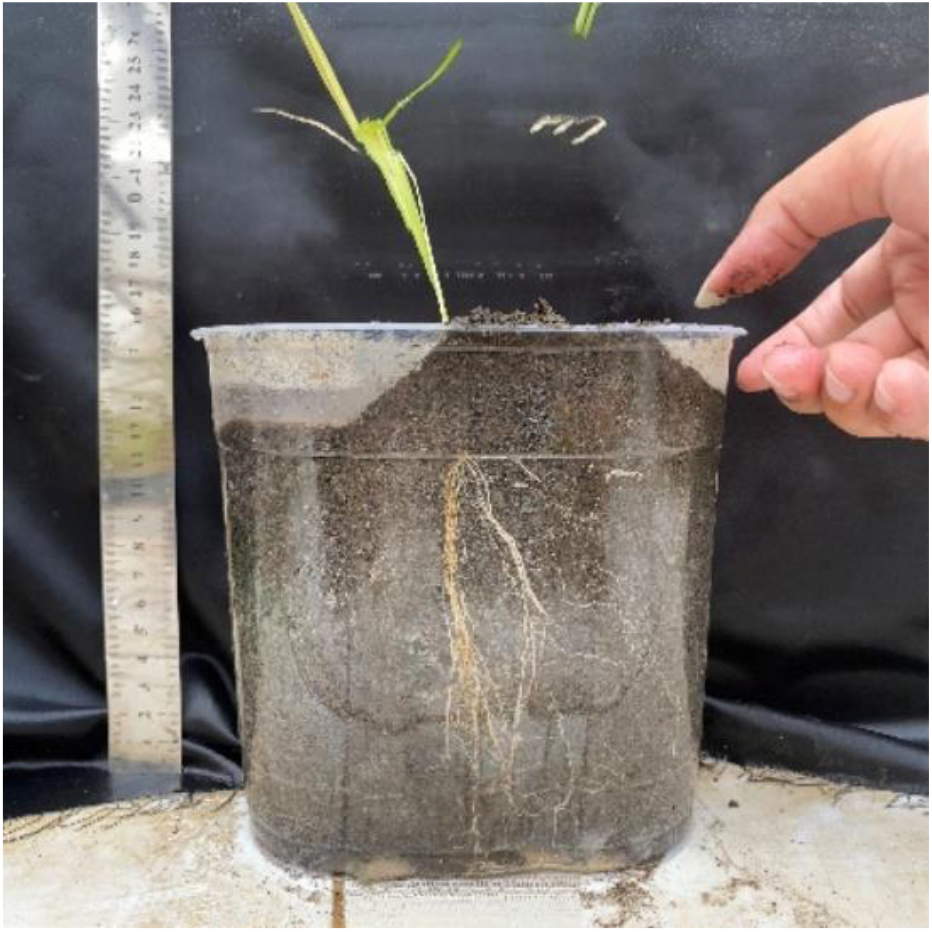
Hasil foto panorama genotipe M70-D

**Gambar 15.**
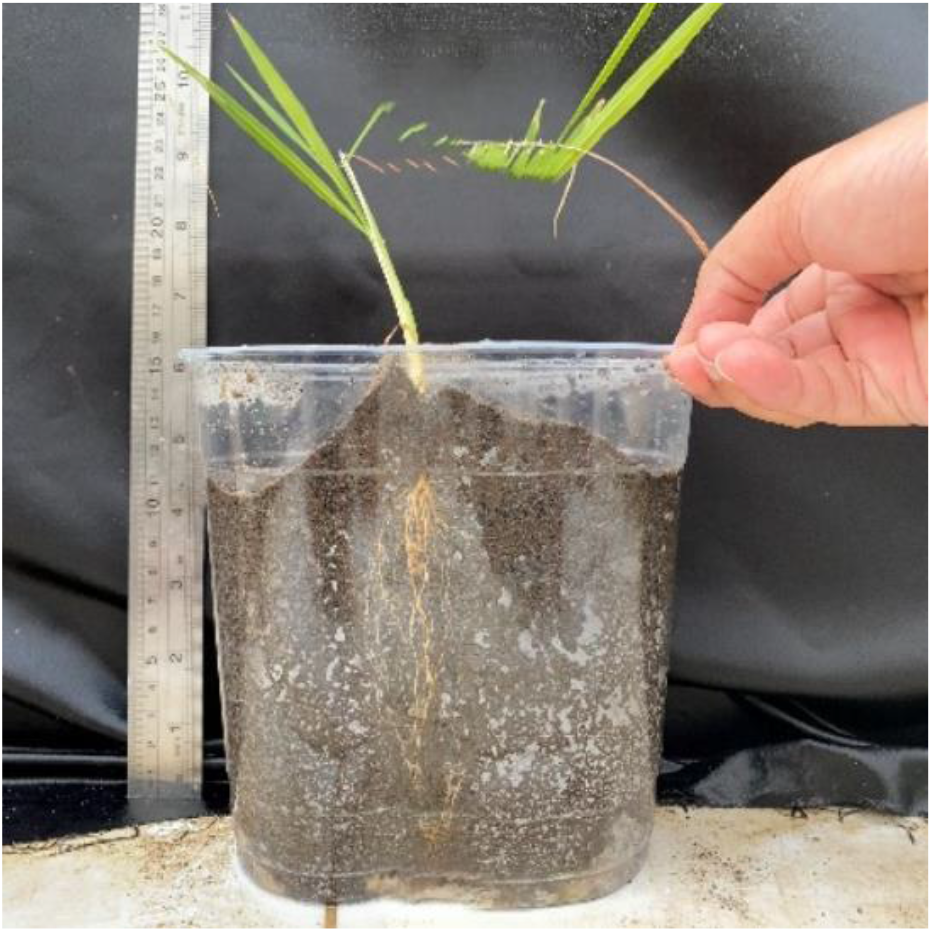
Hasil foto panorama genotipe IR-64

**Gambar 16.**
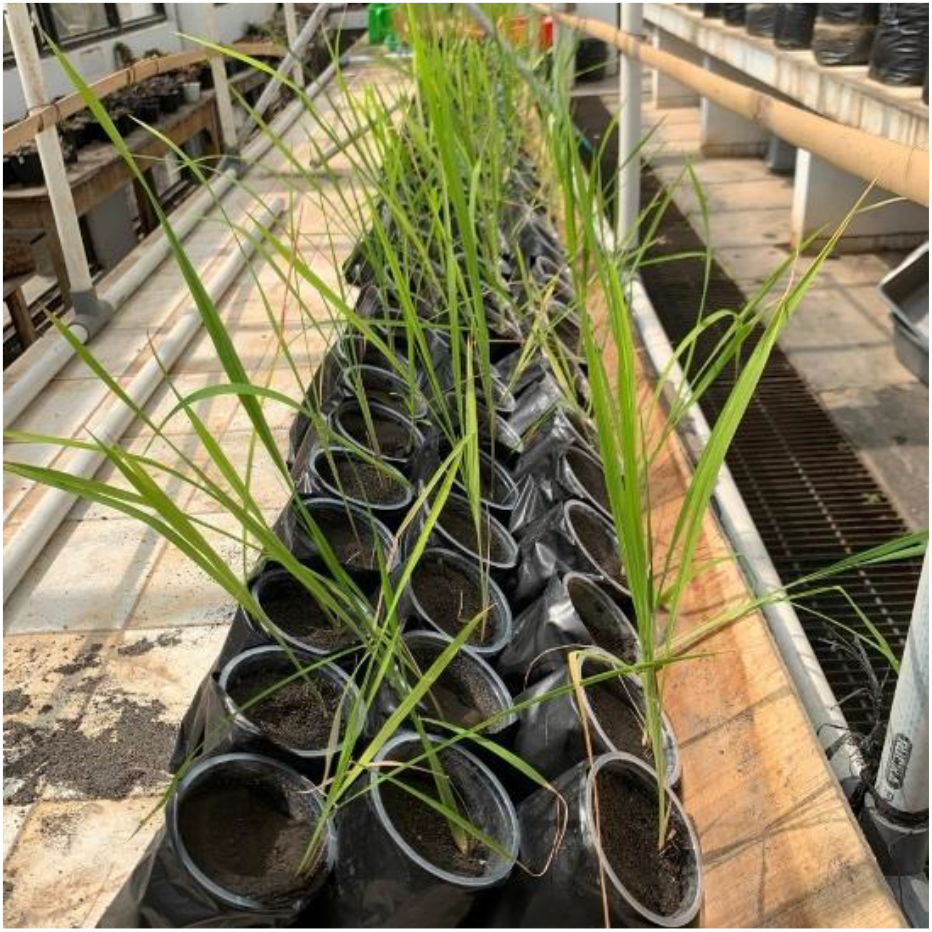
Pengamatan penggulungan dan pengeringan daun

**Gambar 17.**
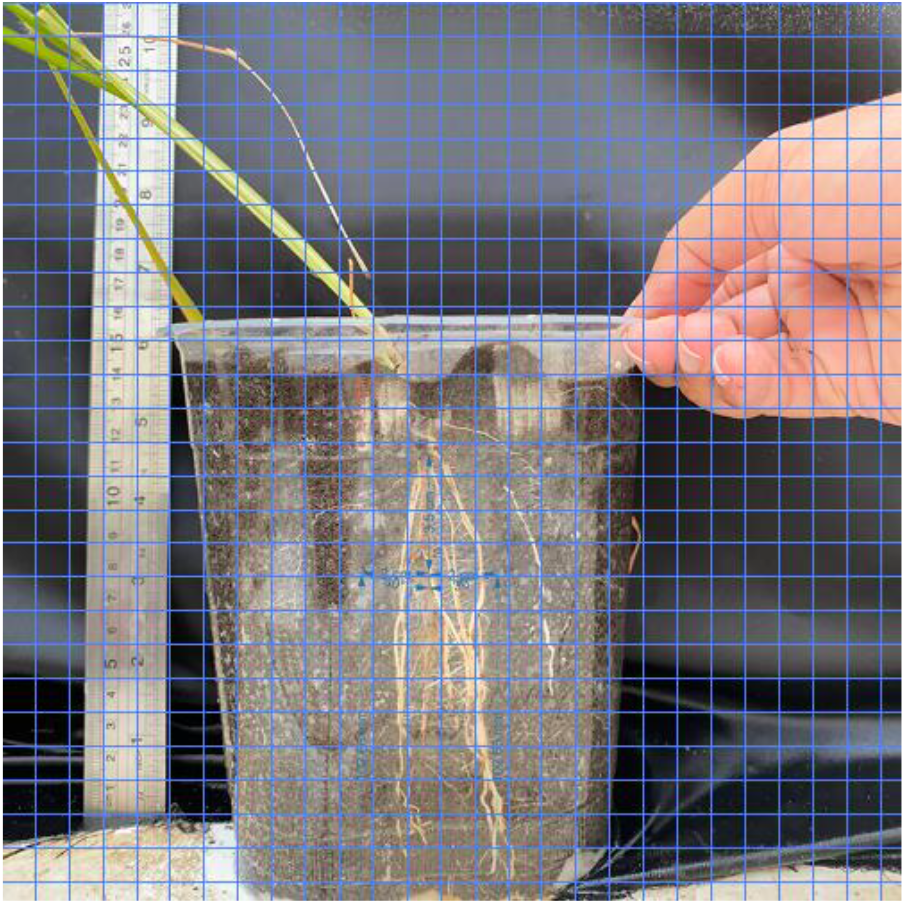
Analisis fraktal pada ukuran frame 1×1cm

**Gambar 18.**
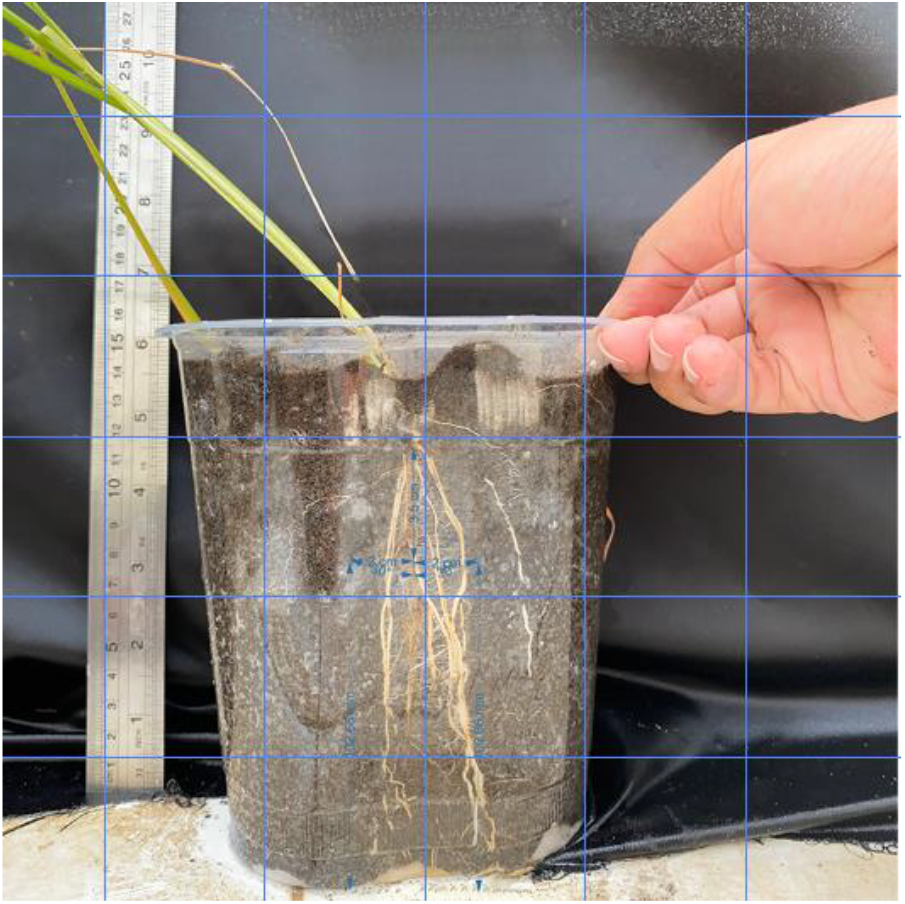
Analisis fraktal pada ukuran frame 5×5cm

**Gambar 19.**
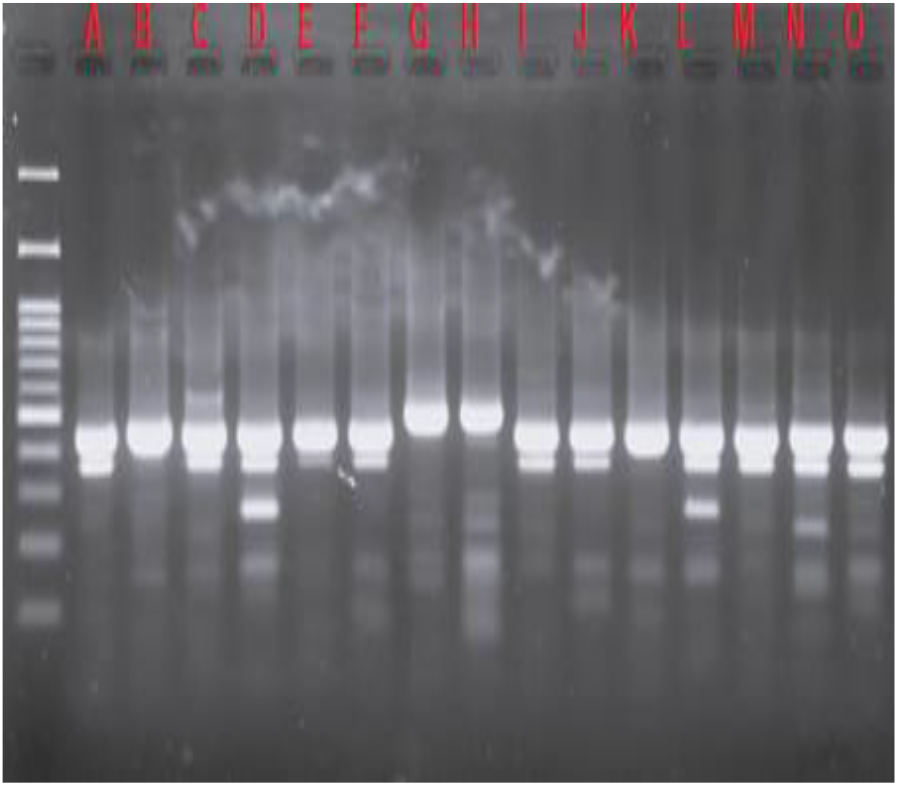
Visualisasi hasil elektoforesis marka DRO-LIR

**Gambar 20.**
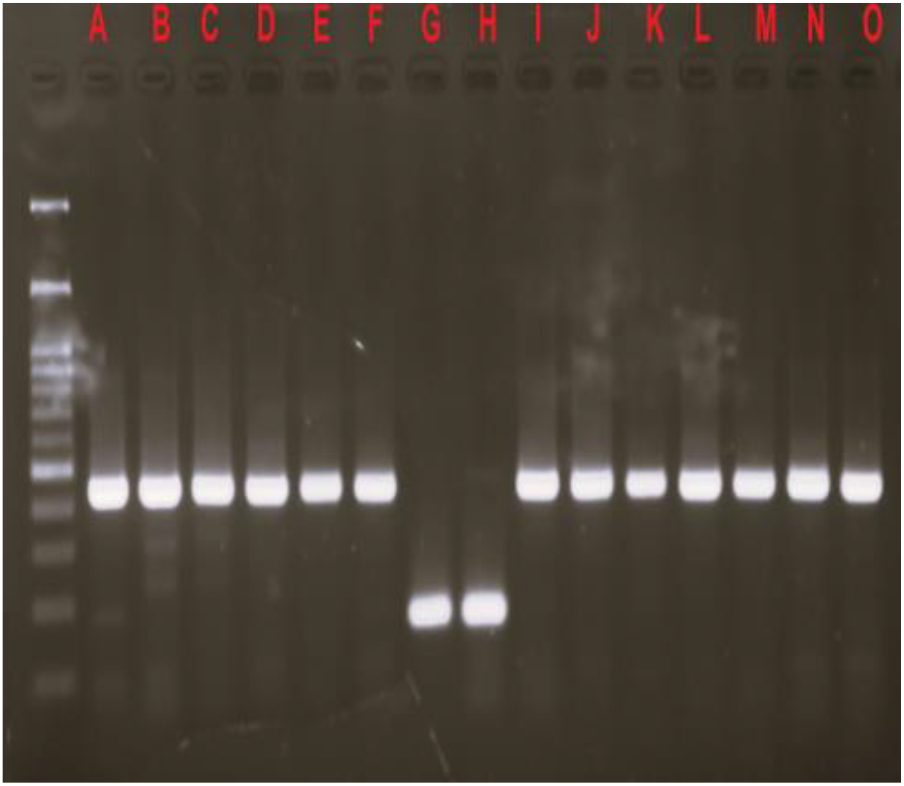
Visualisasi hasil elektoforesis marka DRO-LKP

**Gambar 21.**
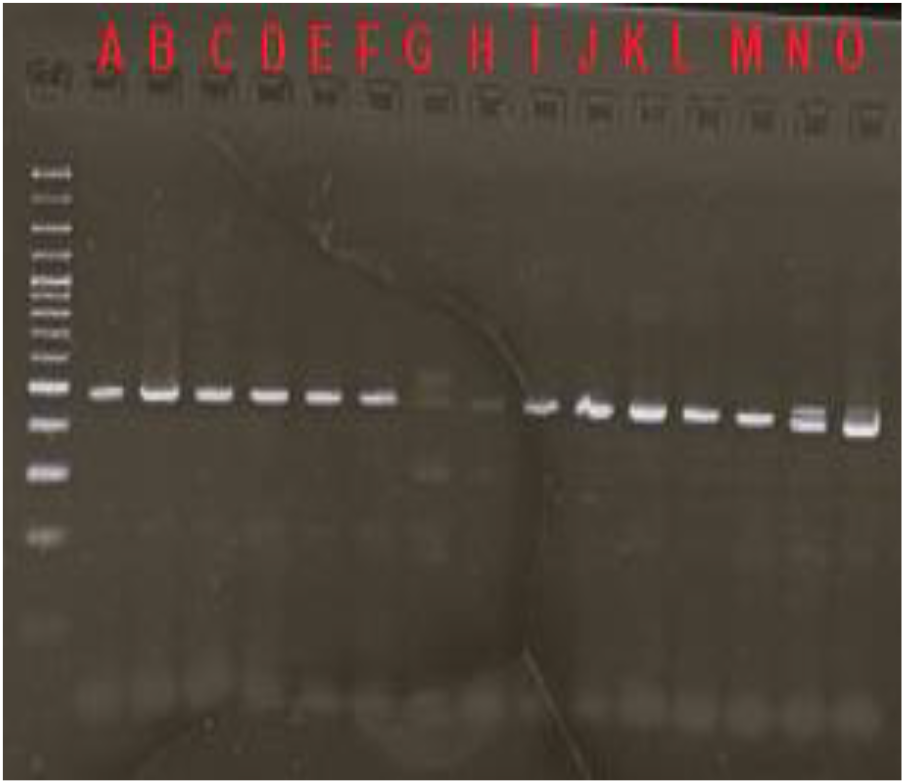
Visualisasi hasil elektoforesis marka RM-7424

**Gambar 21.**
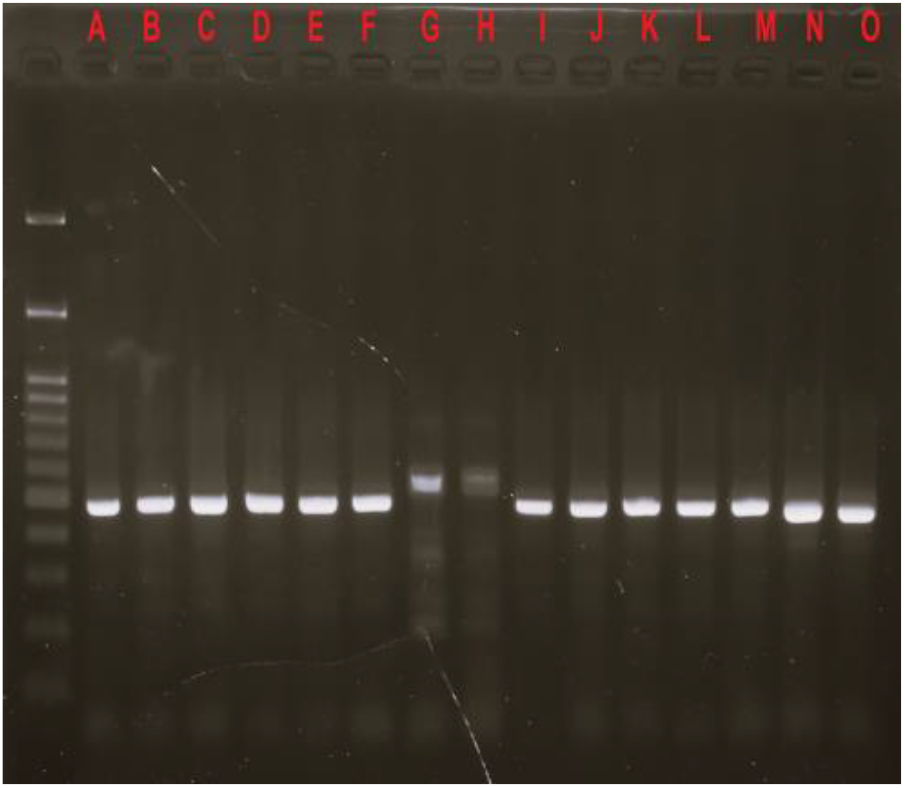
Visualisasi hasil elektoforesis marka RM-24393

**Gambar 21.**
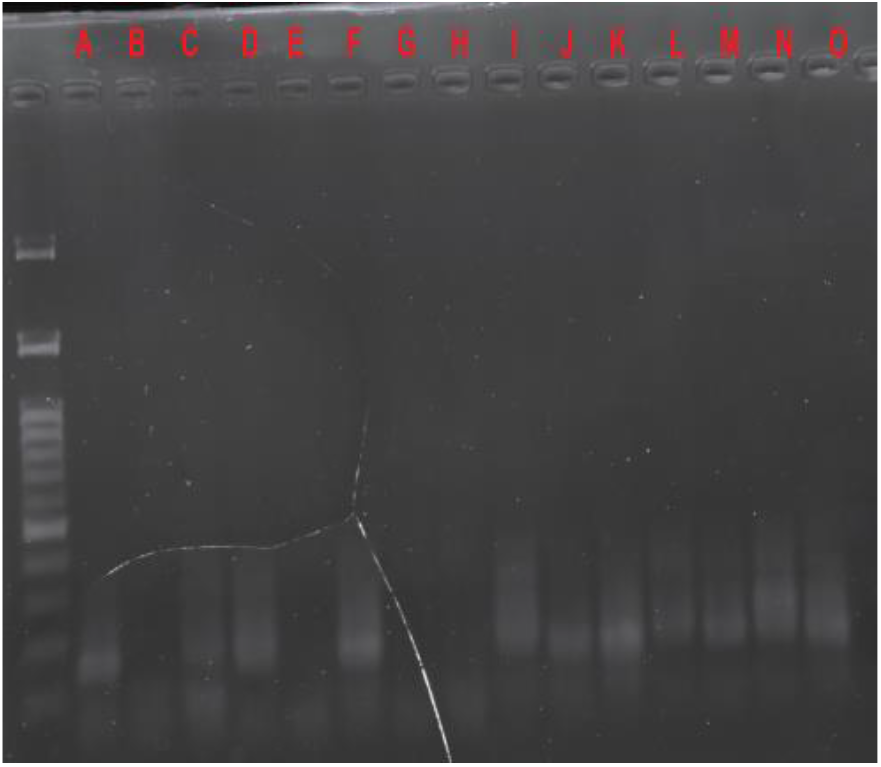
Visualisasi hasil elektoforesis marka RM-72

**Gambar 21.**
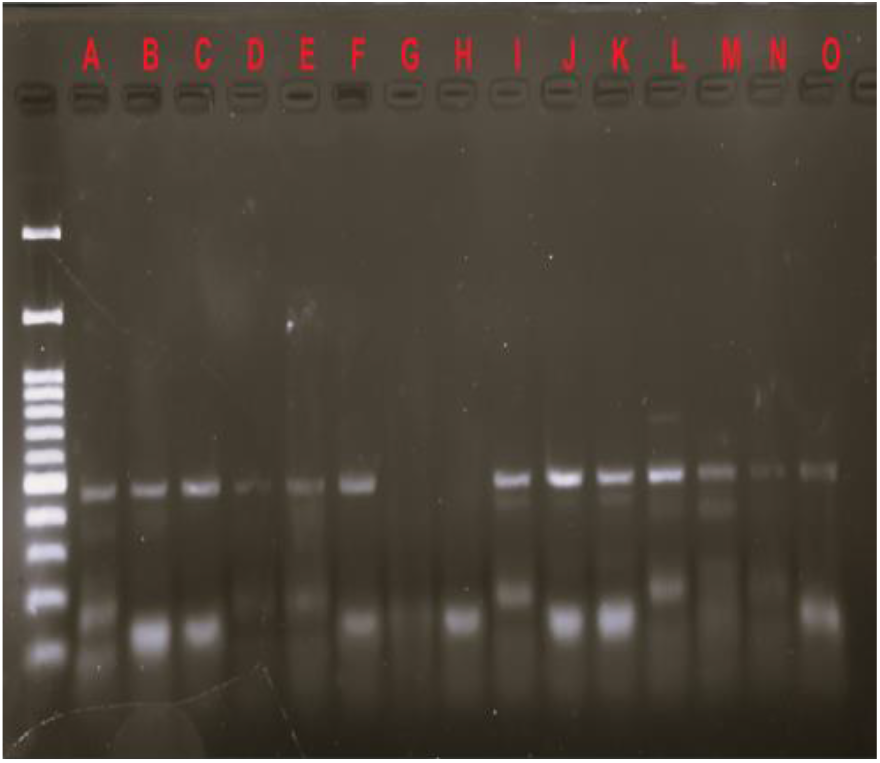
Visualisasi hasil elektoforesis marka RM-228

**Gambar 21.**
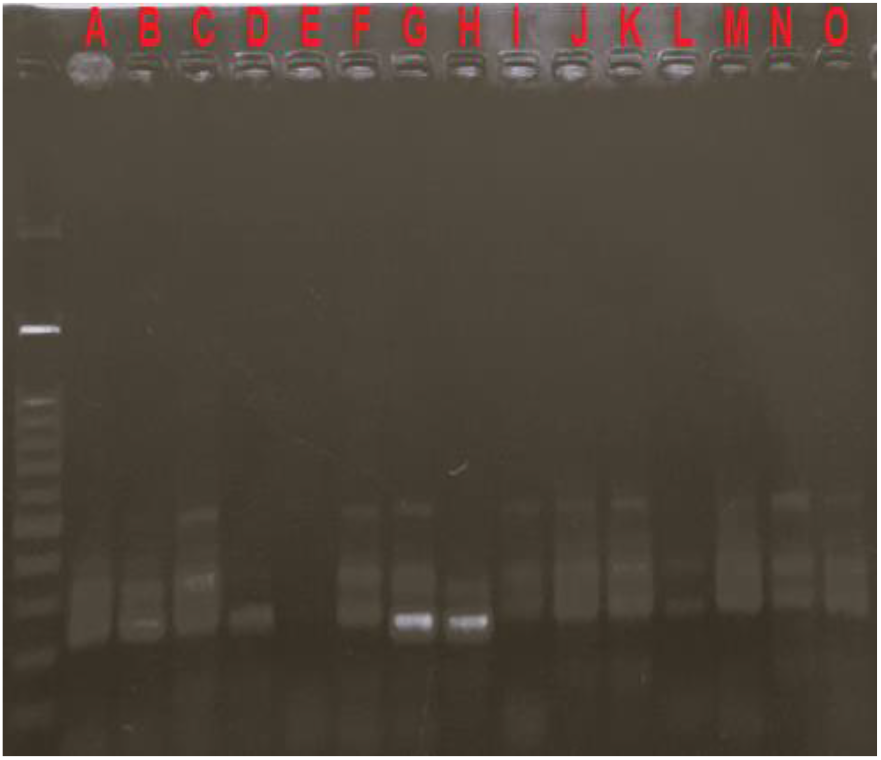
Visualisasi hasil elektoforesis marka RM-518

**Gambar 21.**
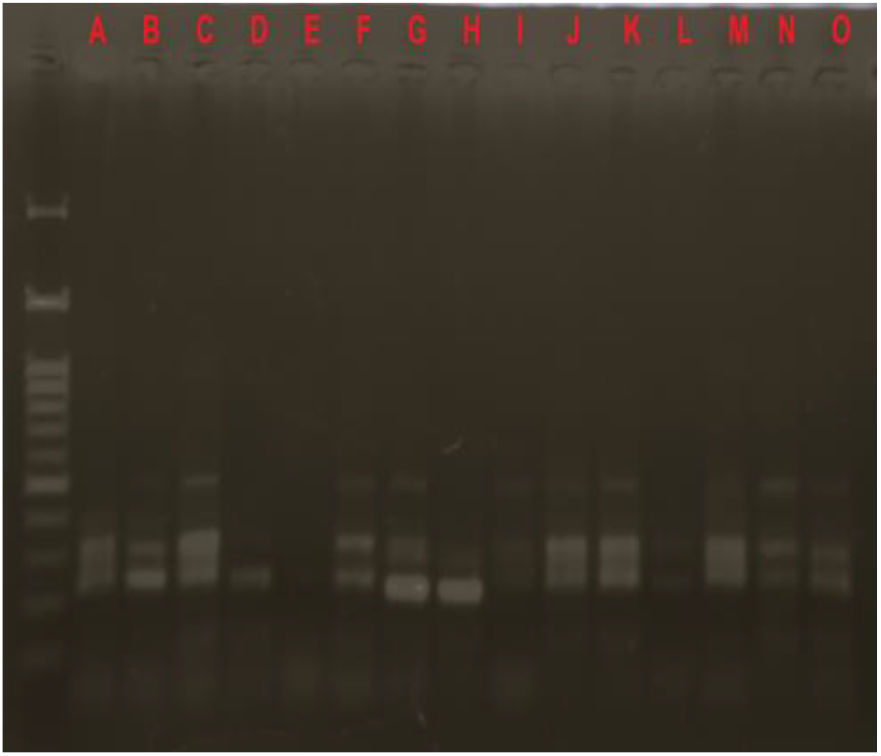
Visualisasi hasil elektoforesis marka RM-20 A

